# Mutation of the *white* gene in *Drosophila* has broad phenotypic and transcriptomic effects

**DOI:** 10.1101/2025.03.03.639943

**Authors:** April Rickle, Krittika Sudhakar, Alix Booms, Ellen Stirtz, Adelheid Lempradl

**Affiliations:** Department of Metabolism and Nutritional Programming, Van Andel Institute, 333 Bostwick Avenue, Grand Rapids, MI 49503, USA

## Abstract

The *white* (*w*) gene, one of the most widely used genetic markers in *Drosophila* research, serves as a standard background mutation for transgene insertions and genetic manipulations^1^. While its primary function involves eye pigmentation, mutations in *white* have been associated with diverse phenotypic effects, including those related to metabolism, behavior, and stress responses^2–19^. However, many studies using these mutants do not account for differences in genetic background, raising concerns about the interpretation of experimental results. To ensure that the observed phenotypic differences are attributable to *white* itself, rather than other genetic differences due to background, we established isogenic fly strains through backcrossing that differ only by the presence or absence of the *white* gene. Given the likely metabolic consequences of *white* gene deletion and its crucial role in neurotransmitter production, we focused our phenotyping assays on behavioral, metabolic, and fitness-related outcomes and performed transcriptomic analysis on adult fly heads. Our findings reveal widespread changes in adult brain gene expression and behavioral, metabolic, and fitness traits, demonstrating that loss of *white* influences multiple biological processes beyond its established role in eye pigmentation. These results emphasize the necessity of genetic background control in *Drosophila* research and warrant caution when using *white* mutants as a baseline for comparative studies.

## Results

First described in 1910 by Thomas Hunt Morgan^20^, the *white* (*w*) gene encodes an ABC class transporter involved in the intracellular transport of eye pigment precursors (tryptophan, guanine, and kynurenine) and other small molecules (GMP, guanine, amine, riboflavin, xanthine, zinc, and pyruvate)^6,9,10,14,21–24^. Its disruption prevents pigment biosynthesis, resulting in white eyes. Due to resource availability and ease of selection, functional *white* (w^+^) has been extensively used as a marker for transgene insertions into *white* mutant (w^-^) backgrounds^1^. Although often considered a “neutral” mutation, numerous studies report phenotypic effects across a wide range of biological processes, including behavior, neurodegeneration, metabolism, lifespan, immunity, and oxidative stress^2–19^. Some of these phenotypes can be attributed to retinal degradation due to pigmentation loss and inability to filter excess light. This deficit has long been associated with abnormalities in behaviors requiring sight^2,6,9,11^. However, *white*’s role in pigmentation cannot explain differing phenotypes such as locomotor degeneration, recovery from anoxia and anesthetics, activity, olfactory learning, cholesterol homeostasis, and lifespan^3,5,6,10–13,15,17,25–28^. Unfortunately, the use of diverse w^-^ fly lines (isoCJ1, w^1118^, w^1^, w^a^, w^a4^, and w^+^) and w^+^ lines (w^e^; Canton-s, Oregon-R, Vallecas, g^1^, and ry^50^) in such publications makes comparisons difficult^3,5,6,10–13,15,17,25–28^. Additionally, due to their short lifespan, small population sizes, and repeated bottleneck events, fly lines maintained separately for many generations are bound to diverge genetically^29–32^. This genetic drift and accumulation of genetic variations over time has been observed in laboratory stocks dating back to 1962^31^. Most of the above studies do not control for genetic background, with only a minority of studies addressing this critical variable^2,4,10,11,14^. A systematic comparison of isogenic w^+^ and w^-^ flies is essential to clarify the broader physiological consequences of this popular genetic marker.

To establish a working isogenic model, we performed ten generations of backcrossing to insert a wild-type copy of *white* from a Harwich line into the commonly used w^1118^ line (BDSC 3605), whose loss of function stems from a partial deletion^33^ of *w*. Heterozygote females from generation 10 were then crossed with w^1118^ males to yield 1:1:1:1 w^+^ males, w^+^ females, w^-^ males, w^-^ females (figure S1). The progeny were then used to determine the effects of w^-^ in otherwise isogenic flies. Age and sex matched flies are used for all experiments and comparisons.

We tested behavioral effects through locomotor, activity, and social spacing assays; metabolic effects through starvation resistance, triglyceride content, and oxidative stress levels; fitness through immune system reactivity, fertility, fecundity, and longevity; and molecular effects through RNA-seq analysis.

### Behavioral phenotyping reveals differences in activity patterns and climbing ability

Despite *white*’s role in the biosynthesis of neurotransmitters, histamine, and melatonin^9^, analysis comparing wake/sleep patterns of w^+^ and w^-^ flies with identical genetic backgrounds at different ages is currently lacking. *Drosophila* circadian rhythms are primarily driven by light, especially in controlled laboratory settings which minimize temperature fluctuations^34^. Aged wild-type and neurodegenerative disease model flies tend to experience changes in activity levels and disturbed sleep compared to younger or healthier counterparts^35,36^. We measured animal movement over time using the Trikinetics Drosophila Activity Monitor (DAM2) in conjunction with the analysis program SCAMP, which calculates 51 different activity and sleep behavior measures^37^. w^-^ flies, while suffering from reduced visual acuity^6^, still exhibit functional circadian rhythms (figures S2E-R). To capture how these variables might contribute to differences between w^+^ and w^-^ flies, we performed principal component analysis (PCA) of all measurements produced with SCAMP (figures 1A and 1B). The first principal component (PC1) separates w^+^ and w^-^ flies and accounts for 35.7% and 41.6% of variability in the male and female dataset, respectively. To determine the variables driving these differences, we isolated and plotted the top 10 measures contributing to PC1 (figures 1C and 1D). This analysis reveals in both sexes and across different ages that w^+^ flies show an increase in sleep-related parameters, while w^-^ flies show an increase in activity and wake parameters (figures S2A-D). While the factors making up PC1 indicate changes in overall behavioral patterns, they cannot fully identify significant differences. For a more traditional statistical approach, we performed 2-way ANOVA using the SCAMP results. This analysis agrees with the PCA results, showing significantly increased sleep-related parameters for w^+^ flies and activity/wake-related parameters for w^-^ flies. In summary, we present the first account of activity and sleep behavior differences in isogenic w^+^ and w^-^ flies. Our results show that while the circadian rhythm remains intact, w^-^ significantly affects sleep/wake patterns, leading to decreased sleep and increased activity levels.

**Figure 1:**
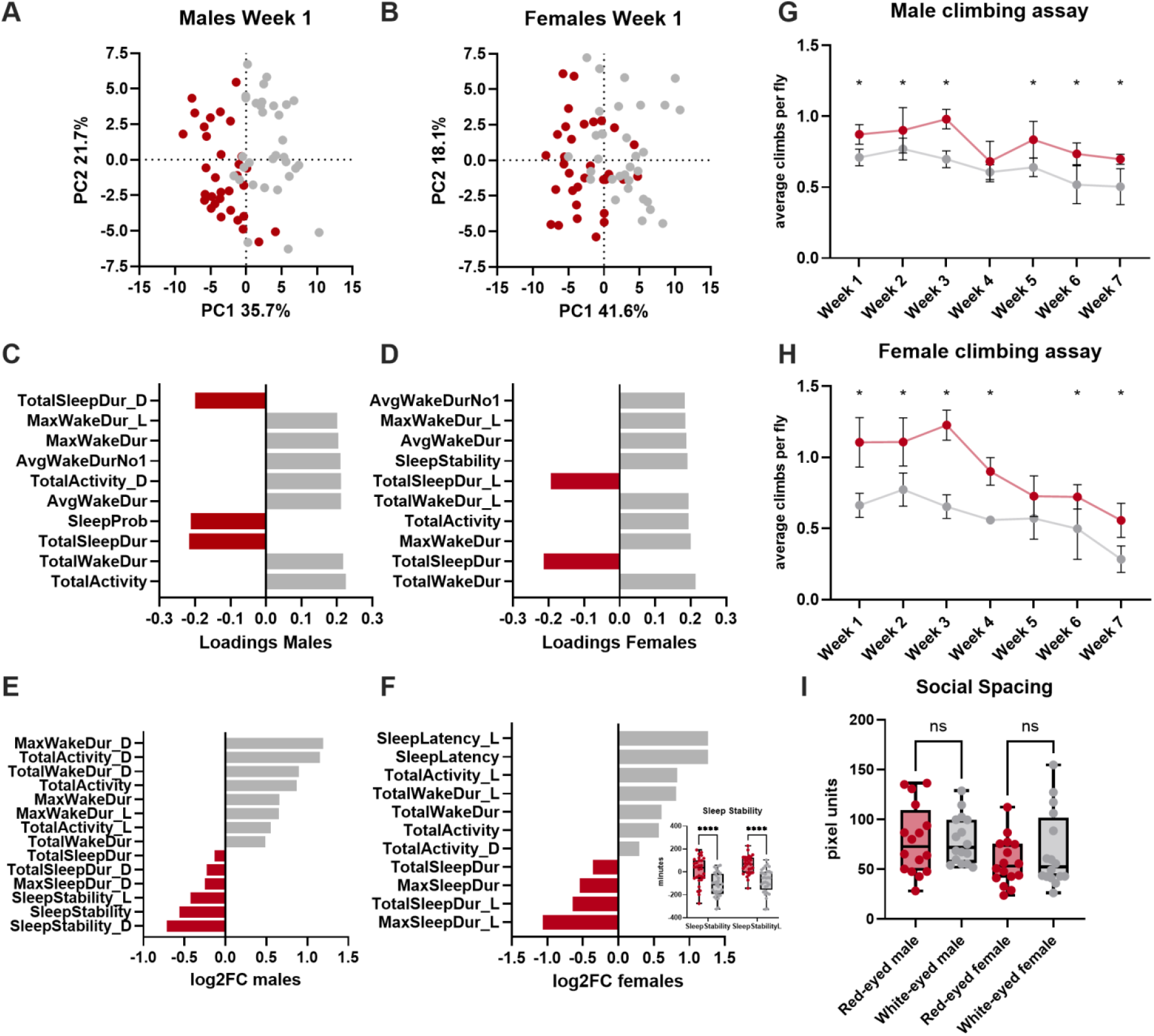
Behavioral phenotyping reveals differences in activity levels and age-related locomotor degeneration but not social spacing. **A and B)** Principal component analysis of activity data analyzed with SCAMP. Each dot represents one fly, and its position in the graph relative to the other dots reflects the similarity in activity patterns between individuals. Both sexes are separated by the first principal component, which accounts for 35.7% and 41.6% of variability in the dataset in males and females, respectively. N=32 for each activity assay. See also figure S2. **C and D)** The top 10 contributing measurements to principal component 1 (PC1) **E and F)** Log2 fold change for significant activity measures identified via ordinary 2-way ANOVA and Šídák’s multiple comparisons test in 1-week old flies. Raw sleep stability values for females are shown in a sub-graph, as the ratios used for log2FC calculations were negative for these measures, making it impossible to calculate a real number. **G)** Average number of climbs per replicate at the 15-second post-shake mark was calculated for 20 total shakes. A full effects model two-way ANOVA indicated an overall significant difference between red and white-eyed flies. N=5 replicates of 10 flies each for each climbing assay. **H)**Average number of climbs per replicate at the 15 second post-shake mark was calculated for 20 total shakes. 2-way ANOVA indicated significance at some weekly timepoints, but not all. **I)** Social spacing. Median distance to nearest neighbor was calculated for each replicate to input into the final dataset. D’Agostino and Pearson testing indicated gaussian distribution and a one-way ANOVA test indicated no significant difference between red and white-eyed flies. N=16 replicates of 20 flies each.

Retinal degeneration is a common measure for neurodegeneration in *Drosophila. white* mutation has been linked to retinal degeneration, which worsens with age^6,13^, suggesting a potential role in age-related neurodegeneration. However, this retinal degradation could simply be due to eye pigment loss and exacerbated over time by light exposure. To assess neurodegeneration independent of eye pigmentation, we measured locomotor ability using the Rapid Iterative Negative Geotaxis (RING) assay^12,18,36^. Previous studies have shown decreased locomotor ability in aged w^-^ flies, though these studies utilized non-isogenic lines and included limited timepoints^2,4,5^. To assess the locomotor ability of our isogenic w^+^ and w^-^ flies throughout their approximate median lifespan, we tested them weekly from 1-7 weeks post-eclosion. At all timepoints, w^-^ flies averaged worse locomotor function than w^+^ flies, with this difference being significant at most weeks (figures 1G and 1H). However, the rate of locomotor deterioration remained similar between w^+^ and w^-^ flies. Therefore, while w^-^ flies have a lower baseline locomotor ability than w^+^ flies, there is no evidence for worsened age-related locomotor decline.

*Drosophila* social behavior is often attributed to scent and pheromones, but can be affected by visual cues as well^38–40^. Differences in social behavior between 4-5 day old w^+^ Canton-s (Cs) and w^-^ w^1118^ flies backcrossed to Cs (w^1118^Cs^10^) have previously been documented, with speculation that visual acuity differences are partly responsible^2^. To determine whether our isogenic w^+^ and w^-^ flies mirrored these results, we performed the social spacing assay and calculated nearest neighbor distances with the cohort. Unexpectedly, we detected no significant differences in social behavior between w^+^ and w^-^ flies (figure 1I). It is possible that while the Cs and w^1118^Cs^10^ lines used in the published study were initially isogenic, maintenance as separate stocks for many generations allowed for genetic drift over time. These results imply that establishment of social spacing does not involve visual acuity and emphasize the confounding role of laboratory stock genetic drift in behavioral assays.

### Metabolic phenotyping reveals sex specific differences in triglyceride levels and starvation resistance

While *white* is not involved in tryptophan uptake from the extracellular space, its role in intracellular tryptophan transport is interconnected to downstream metabolism^41,42^. Previous experiments show that w^-^ flies have altered levels of metabolites from the kynurenine pathway (tryptophan, kynurenine, kynurenic acid, 3-hydroxykynurenine, and xanthurenic acid), purine metabolism (guanine, guanosine, xanthine, and urate), folic acid, vitamin B2^14^, serotonin, dopamine, and histamine^9^. Tryptophan metabolism is also linked to lipid metabolism in mice and human tissue culture, and cholesterol homeostasis in flies^10,43^. Increased dietary tryptophan intake in rats is associated with reduced serum triglyceride levels, and increased tryptophan levels are associated with increased lipid peroxidation, a common readout for oxidative stress^44,45^.

Due to tryptophan’s important role in whole body metabolism, we measured the effects of *w* mutation on metabolic phenotypes. First, we compared body weight and triglyceride content of w^+^ and w^-^ flies. While body weight is not significantly different (figure 2A), w^+^ males show significantly higher triglyceride content than w^-^ males (figure 2B), with no significant differences observed in females.

**Figure 2:**
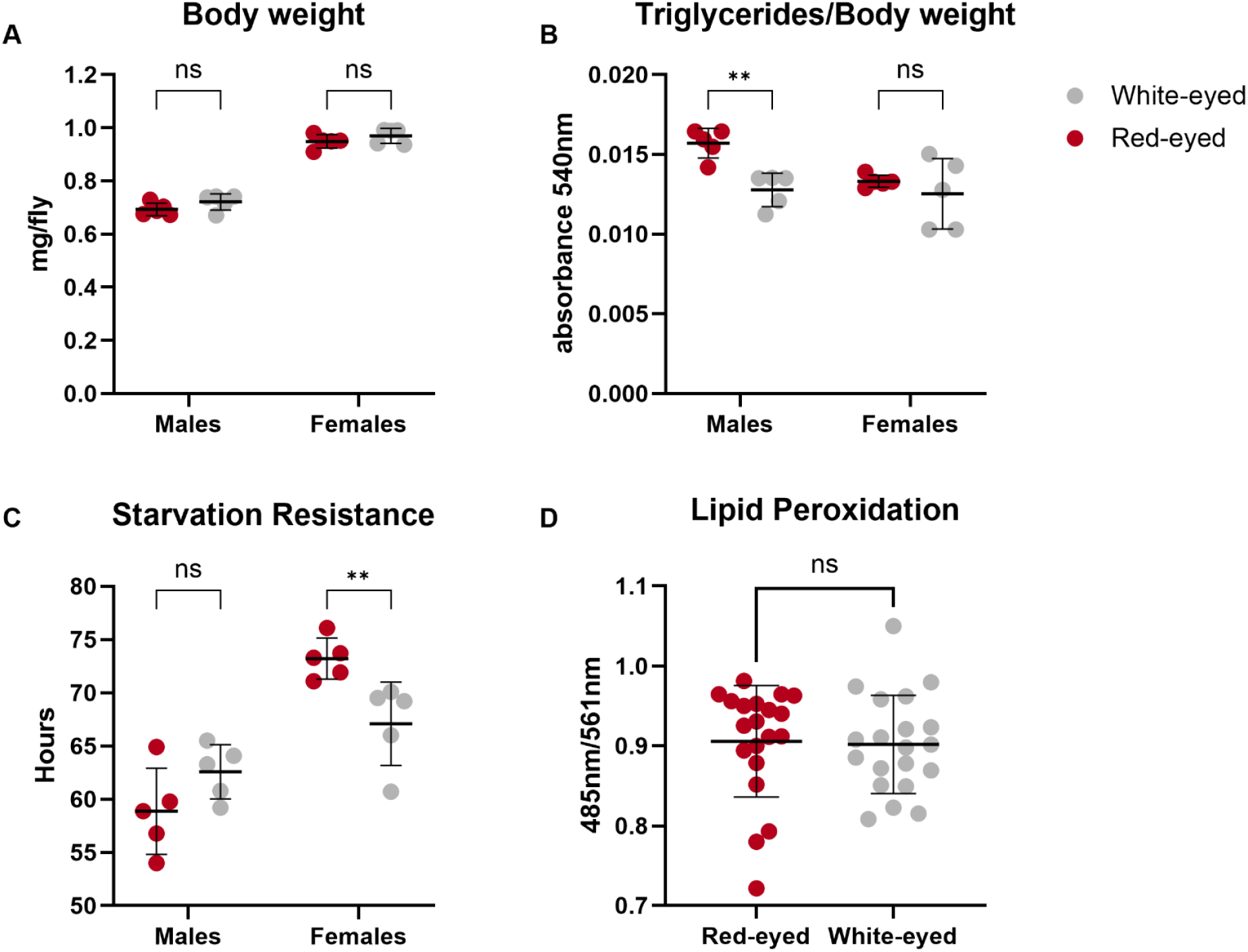
Mutation of the *white* gene affects triglyceride levels and starvation resistance in a sex-specific manner. **A)** No significant difference in body weight per fly was observed between w^+^ and w^-^ males and females. Statistics were calculated via ordinary 2-way ANOVA with Šídák’s multiple comparisons test. N= 5 replicates of 10 flies each for the triglyceride and body weight assays. **B)** Measuring triglyceride levels reveals that w^-^ males have significantly lower triglyceride content per body weight of fly compared to w^+^ males, while there is no significant difference observed in females. Statistics were calculated via ordinary 2-way ANOVA with an uncorrected Fisher’s LSD. **C)** Measuring starvation resistance revealed that w^+^ females have a significantly higher resistance to starvation compared to w^-^ females, while there is no significant difference observed in males. Statistics were calculated via ordinary 2-way ANOVA with an uncorrected Fisher’s LSD. N= 5 replicates of 10 flies each. **D)** No significant difference was found between homozygous w^+^ and w^-^ embryos when measuring lipid peroxidation. Statistics were calculated via Mann-Whitney test. N= 20 replicates of 10 individuals each.

As triglyceride content correlates with starvation resistance, we performed a starvation resistance assay^46^. A previous study using non-isogenic strains reported that w^-^ flies have lower starvation resistance than w^+^ flies^6^. Our results show no significant differences in starvation resistance for males despite having significant changes in triglyceride content. Interestingly, w^-^ females show decreased starvation resistance compared to w^+^ females, despite not showing any significant difference in triglyceride content (figure 2C). These results indicate sex-specific effects of w^-^ on metabolic phenotypes and partially validate previous findings^6^.

Oxidative stress is linked to neurodegeneration in both humans and flies, and w^-^ flies have lower resistance to oxidative stress from paraquat or H^2^O^2^ exposure^6^. These findings suggest a link between *w* mutation and oxidative stress. To test whether *w* affects oxidative stress, we measured lipid peroxidation levels in embryos. As w^+^ and w^-^ embryos are indistinguishable, we generated homozygous w^+^ and w^-^ embryos from our isogenic crosses for these experiments. We saw no significant differences in lipid peroxidation between w^+^ and w^-^ embryos (figure 2D).

### Fitness phenotyping reveals a difference in fertility

Egg production requires both energy and nutrients, with protein availability being particularly important^51^. Despite the role of white in tryptophan transport, its effects on egg-laying have not been studied. To assess fecundity and fertility, we counted the number of eggs laid by unmated and mated females, respectively. We observed no significant difference in fecundity; however, fertility is significantly affected with w^-^ flies laying more eggs than w^+^ flies at 21 and 31 days old (figures 3A and 3B), despite w^-^ males’ reduced heterosexual copulation success^6,16,18^.

**Figure 3:**
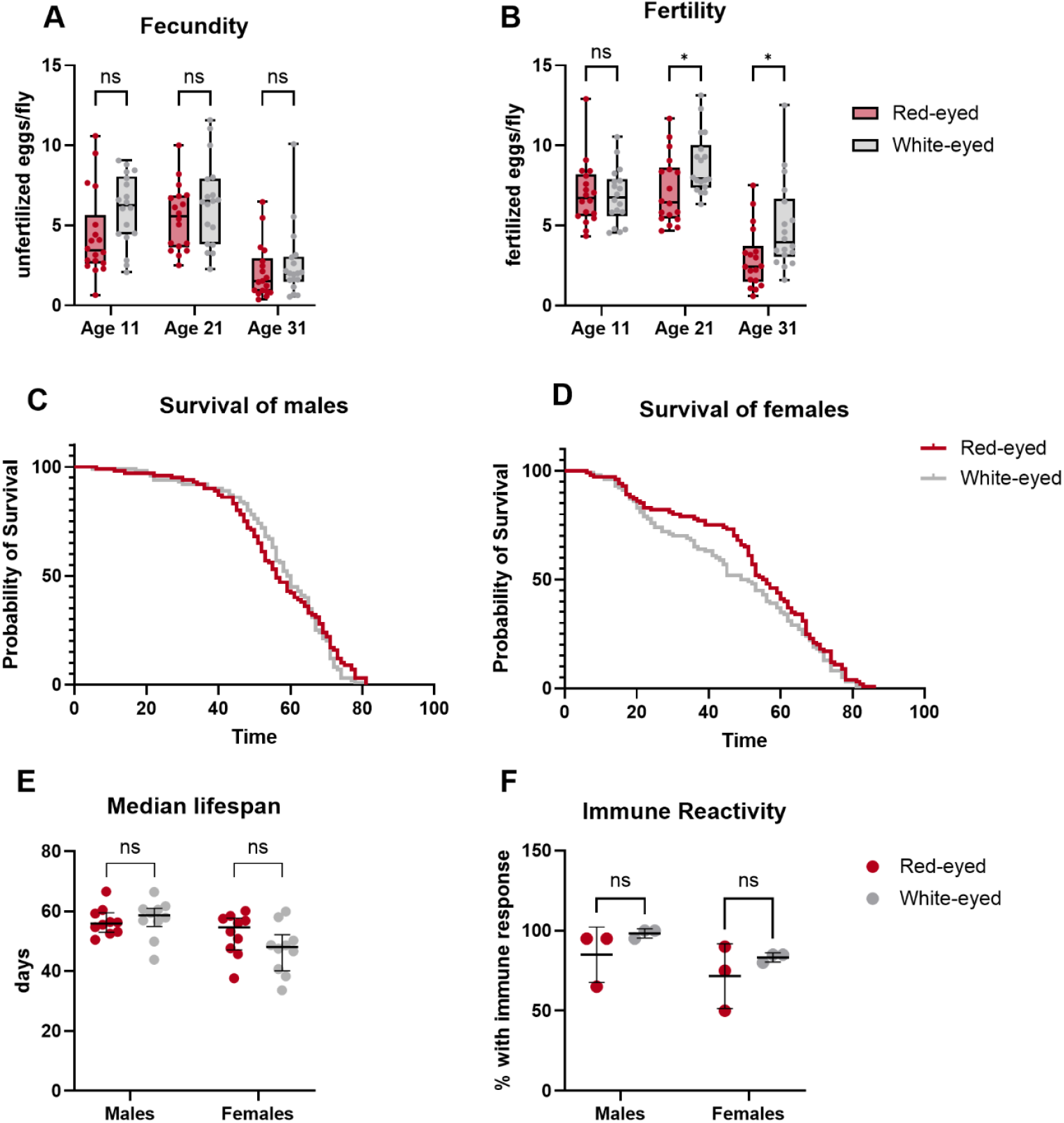
The *white* gene affects fertility. **A)** Fecundity assay (eggs laid per unmated female) shows that there is no significant difference in egg laying between w^+^ and w^-^ females at 11, 21, and 31 days of age. **B)** Fertility assay (eggs laid per mated female) shows that 21 and 31-day old w^-^ females lay significantly more eggs compared to w^+^ females. Statistics for fecundity and fertility were calculated via 2-way ANOVA with Geisser-Greenhouse correction and Tukey’s multiple comparisons test. N=6 replicates of 20 females each, tested across 3 days for the fecundity and fertility assays. **C and D)** Survival curve for males and females, respectively. Neither Mantel-Cox nor Gehan-Breslow-Wilcoxon testing shows a significant difference in the males or female w^+^ and w^-^ curves. **E)** The median lifespan of red and white-eyed males and females shows no significant difference between w^+^ and w^-^ males and females. Statistics were calculated via ordinary 2-way ANOVA with an uncorrected Fisher’s LSD. N=10 vials of 10 flies each for the lifespan assay. **F)** Immune reactivity assay shows no significant difference between w^+^ and w^-^ larvae. Statistics were calculated via 2-way ANOVA with Šídák’s multiple comparisons test. N= 3 replicates of 20 larvae each.

Previous studies in non-isogenic backgrounds suggest that w^-^ lines have different lifespans compared to w^+^ lines^5,6,28^. Notably, some recorded differences contradict each other, pointing towards genetic background as a contributing factor. For example, w^-^ flies have been reported to live longer than Oregon-R, but no difference was shown compared to Canton-s flies, while Oregon-R flies have been shown to live significantly longer than Canton-s flies^5,28,47^. Given this discrepancy, we measured lifespan in our isogenic w^+^ and w^-^ flies. Notably, we observed no significant difference in median lifespan (figures 3C-E), indicating that the previously observed changes were due to genetic background rather than the *white* mutation.

In humans, extensive research indicates a link between tryptophan metabolism, inflammation, and the immune system^48^. A link has also been suggested in *Drosophila*, where metabolites of the kynurenine pathway were shown to interact with zinc and influence the immune system^49^. Due to the link between tryptophan metabolism and the immune system, we performed an immune reactivity assay on w^+^ and w^-^ larvae, but we observed no significant difference (figure 3F).

### RNA-seq reveals general and sex-specific gene expression changes

Given the potential far-reaching effects of aberrant tryptophan metabolism and the observed differences in behavior, we performed gene expression analysis on heads from 1-week old w^+^ and w^-^ male and female flies to assess the system-wide effects of *white* mutation.

Differential gene expression analysis revealed 1,312 and 1,752 total differentially expressed (DE) transcripts in males and females, respectively, with more genes downregulated than upregulated in w^-^ flies of both sexes. As expected, in both males and females, we detected a >99% decrease in *white* expression in w^-^ flies (figures 4A, 4B, and S3A). To better interpret the other gene expression changes, we performed Gene Set Enrichment Analysis (GSEA). While females had a greater number of enriched pathways, the overall results for males and females were similar, with an overlap of many gene sets related to ribosome/translation, glucose and fatty acid metabolism, mitochondria, cell adhesion, and chitin (figures 4C and S3B, tables 1-4). Due to the sex-specific effects seen in our phenotyping analysis, we examined sex-specific gene expression patterns. Despite a significant overlap, there were also 366 and 806 DE genes identified in only males or females, respectively (figure 4D). We performed overrepresentation analysis (ORA) on these sex-specific DE genes to identify related transcriptional programs. Male-specific upregulated genes were mostly related to response to environmental stimuli, while the top 10 male-specific downregulated genes mainly involved carbohydrate and nucleotide metabolism (figures 4E and 4F, tables 5 and 6). The top 10 female-specific upregulated genes were enriched for development-related pathways, while the female-specific downregulated genes involved ribosomes, small noncoding RNAs, and small molecule metabolism (figures S3C and S3D, tables 7 and 8). Our transcriptomic analysis reveals vast differences in pathways between w^-^ and w^+^ flies, as well as additional sex-specific differences, further underlining our phenotyping results.

**Figure 4:**
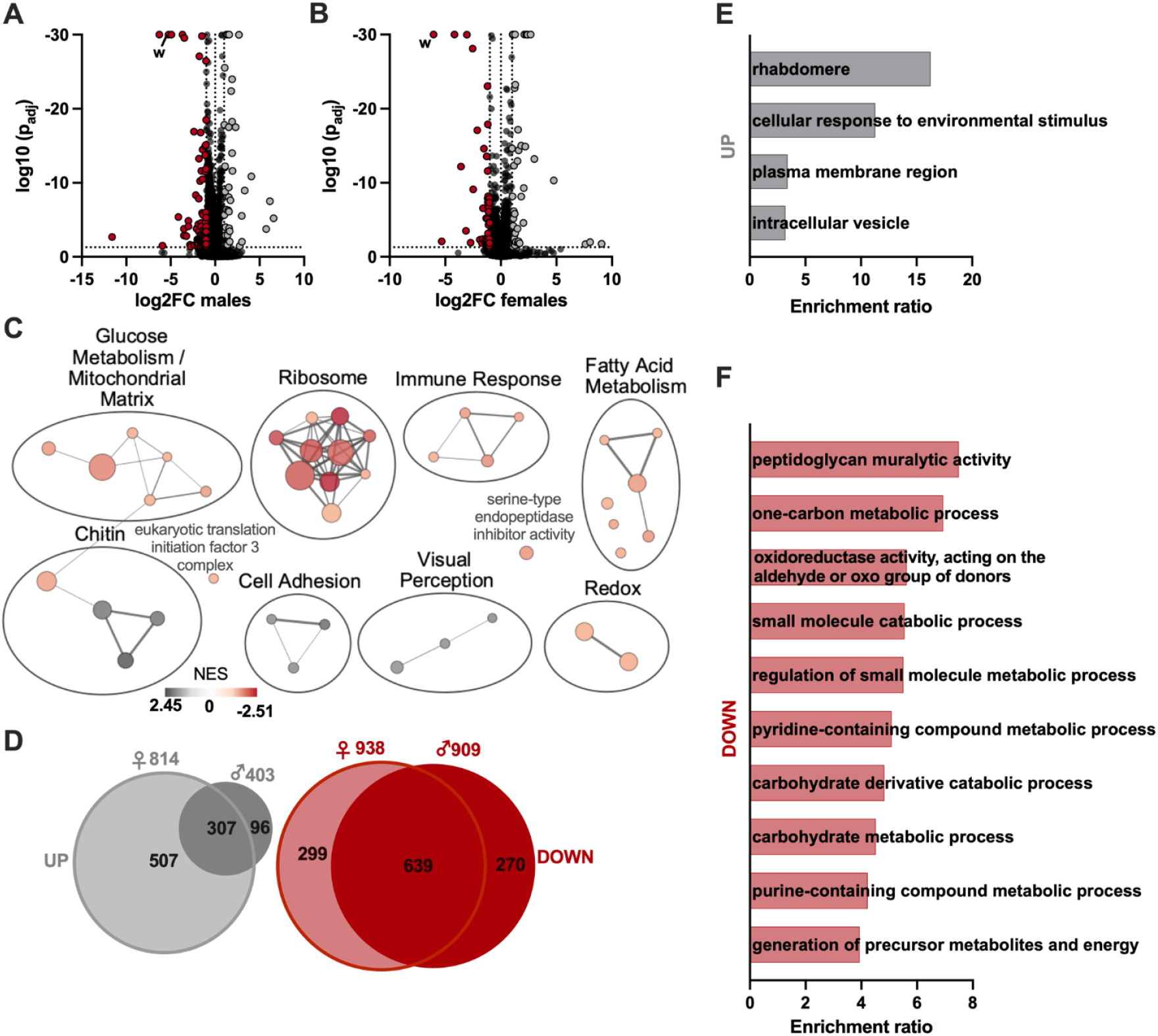
RNA-seq reveals significant both broad and sex-specific gene expression changes. **A and B)** Volcano plots of differentially expressed (DE) genes for males (A) and females (B). Expression of *w* is significantly lower in both male and female w^-^ flies. See also figure S3A. **C)** Gene Set Enrichment Analysis (GSEA) of all DE genes in males. See also figure S3B. **D)** Differentially upregulated (grey) and downregulated (red) genes in females (left circle) and males (right circle). The number above each circle is the total number of up or downregulated genes for that condition, with the numbers inside the Venn diagram showing how many of those genes are unique or shared between males and females **E)** Overrepresentation analysis for upregulated male-specific DE genes. See also figure S3C. **F)** Overrepresentation analysis for the top 10 downregulated male-specific DE genes. See also figure S3D.

## Discussion

*White* is a widely used phenotypic marker in *Drosophila*, with the assumption that mutations have minimal side effects. Using otherwise isogenic flies, we show that this mutation has vast effects on transcription, metabolism, and behavior. While our findings replicate some published results, they also diverge from previous studies assessing *white* function using genetically divergent lines, highlighting genetic drift as a major confounding factor. This work also presents a rigorously controlled transcriptomic dataset isolating the effects of *white* gene mutation, addressing a critical gap in the field.

*Drosophila* are widely used as a model for neurodegeneration, metabolism, and aging; knowing the exact effects of w^-^ is crucial for interpreting disease models built on this background. As science progresses, improved methods allow for detection of more subtle phenotypes, but may also detect deeper effects of “surface level” mutations which have historically been used as markers out of convenience. Here, we employed widely used, sensitive, state-of-the-art phenotyping assays to establish a baseline behavioral assessment of w^-^. The results will guide researchers in interpreting their findings and in selecting or modifying the use of genetic backgrounds.

Our results indicate genetic background as a major driver of phenotypic differences. The *Drosophila* community commonly compares phenotypes in lines with different genetic backgrounds; our results underline the importance of including proper controls for genetic background such as deficiency strains, multiple independently derived alleles of the same gene, rescue experiments, phenotypic comparison of F1 hybrids, and backcrossing into isogenic backgrounds. Efforts should also be made to reduce genetic drift within lab stocks, including increased strain tracking, avoidance of extensive bottlenecking during husbandry, and regular exchange of individuals between duplicate stock vials.

*White* is only one example; similar effects could exist for other popular genetic markers. For example, intracellular transporters scarlet and brown heterodimerize with white, and their mutations likely share some phenotypic and molecular effects with *w* mutants^6^. The curly wing phenotype is caused by mutation of *duox*, a reactive oxygen species generating NADPH oxidase which plays a role in the immune system^50^. Stubble, a transmembrane serine protease, plays an important role in shaping the cytoskeleton in morphogenesis; homozygous mutations are lethal in larvae, indicating its crucial role in *Drosophila* biology^51^.

The issue of genetic drift is not unique to *Drosophila*. The mouse line C57BL/6 (commonly called black 6) has many sub-strains, each with phenotypic differences due to hundreds of generations isolated in various labs and repositories. The Jackson Laboratory, originator of black 6, considers 20 generations of separation sufficient to create a sub-strain^52–54^. Assuming a generational time of 14 days (ie, flipping every 2 weeks), *Drosophila* produce up to 26 generations per year. According to this estimate, the *white* mutant isolated by Morgan in 1910 (w^1^) has spent nearly 3,000 generations as laboratory stock. Over the years, it has also been distributed to countless individual labs and stock centers. As a result, the potential for genotypic and phenotypic differences between different stocks of the same original fly line is substantial, which helps explain phenotypic discrepancies reported in the literature.

## Resource availability Lead contact

Requests for further information and resources should be directed to and will be fulfilled by the lead contact, Adelheid Lempradl

## Materials availability

Strains are available from the corresponding author upon request.

## Data and code availability

The raw RNA-seq data is available on Gene Expression Omnibus under the accession code GSE290227.

The code for RNA-seq analysis is available in the supplemental files.

Any additional information to reanalyze the reported data is available from the lead contact upon request.

## Acknowledgements

The authors are grateful to the Genomics Core at Van Andel Institute for their work performing the RNA sequencing, as well as Joe Roy and Dmitri Martirosov for their work as lab support staff. This research was funded by the Van Andel Institute.

## Author contributions

Conceived and designed the experiments: AR AL. Performed the experiments: AR SK ES. Analyzed the data: AR SK ES AB AL. Wrote the paper: AR AL.

## Main text figure/table legends

**Figure S1:**
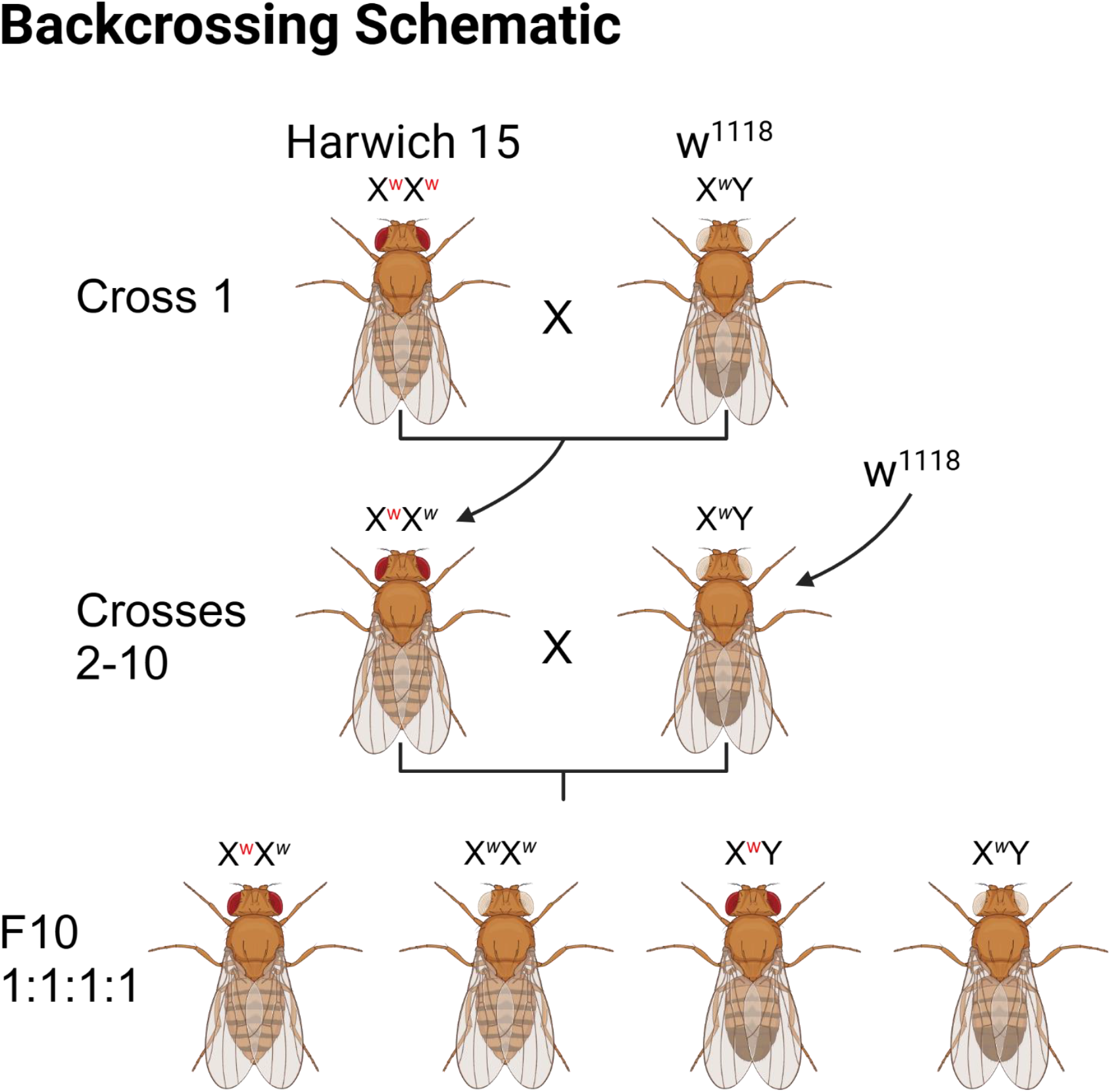
Backcrossing schematic. Cross 1 was between a Harwich 15 female and a w^1118^ male. Virgin female offspring were collected and mated to w^1118^ males for cross 2. Crosses 3-10 took red-eyed females from the previous cross and mated them to w^1118^ males. Flies were collected for assays from cross 10. Flies from cross 10 were also crossed to one another to produce homozygous white-eyed and red-eyed lines for experiments requiring embryos and larvae, which cannot be phenotyped by eye color.

**Figure S2:**
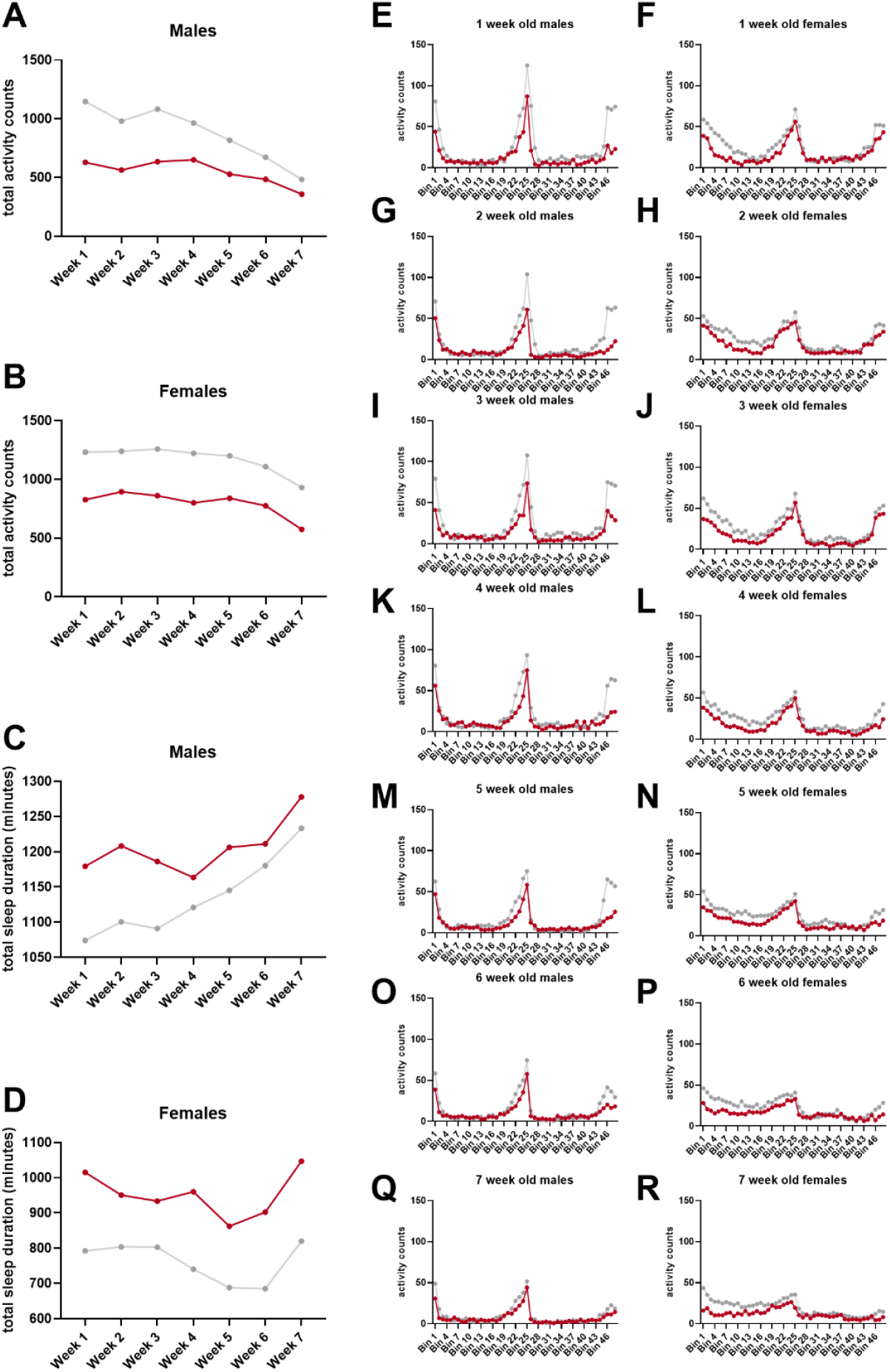
Additional activity assay data shows decreased activity and increased sleep over time in both sexes, related to figure 1. **A and B)** Total activity counts for males and females over 7 weeks **C and D)** Total sleep duration for males and females over 7 weeks **E-R)** total activity counts shown in half-hour bins for 1 week old (E, F), 2 week old (G, H), 3 week old (I, J), 4 week old (K, L), 5 week old (M, N), 6 week old (O, P), and 7 week old (Q, R) males (left column) and females (right column)

**Figure S3:**
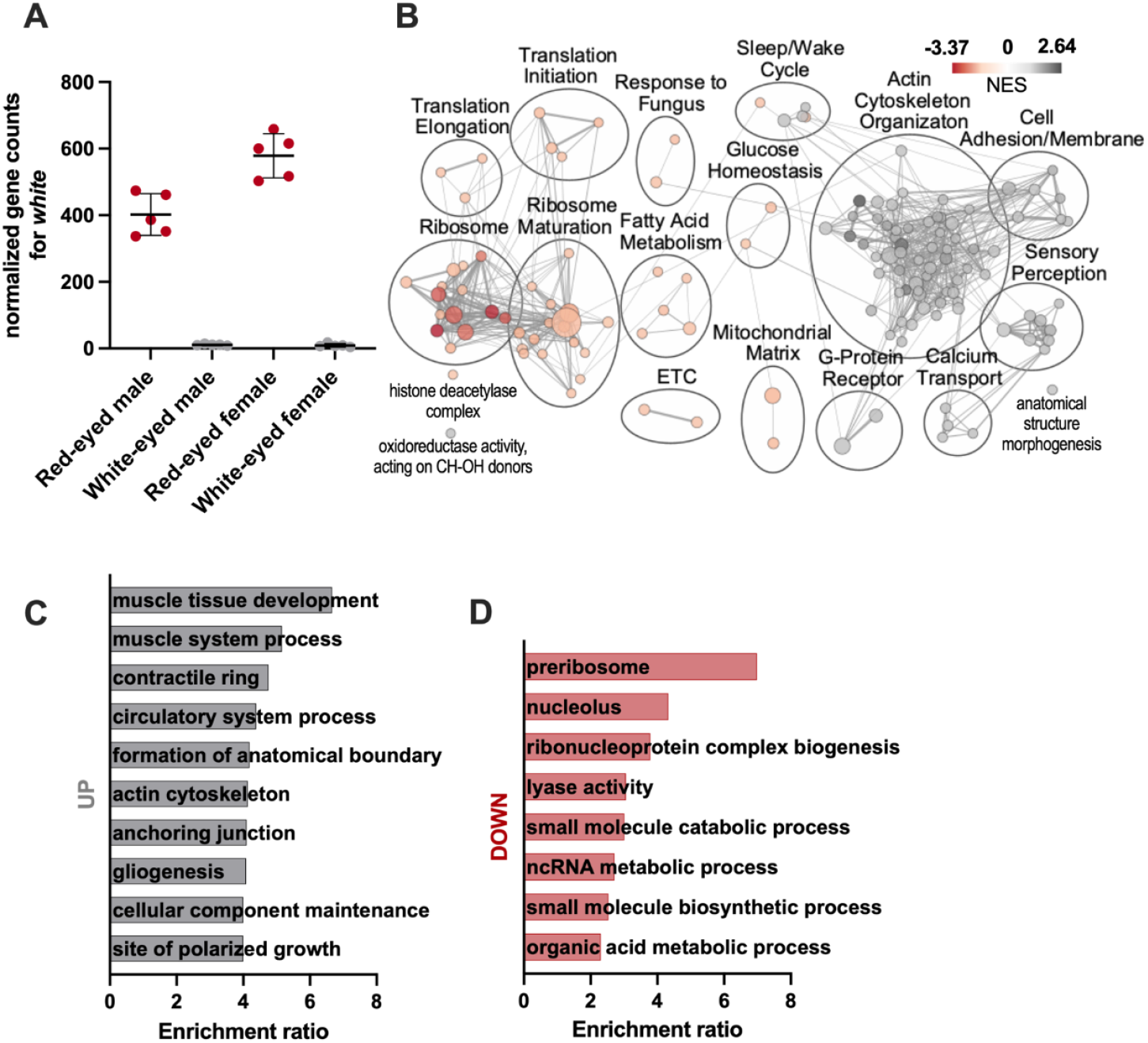
RNA-seq confirms loss of *w* expression, female-specific gene expression changes, related to figure 4. **A)** Expression of *w* in male and female w^+^ and w^-^ flies **B)** GSEA of all DE genes in females **C)** Overrepresentation analysis for the top 10 upregulated female-specific DE genes **A)** Overrepresentation analysis for downregulated female-specific DE genes

## Methods

### Fly husbandry and lines used

Flies were maintained at 25°C at 60% humidity on a 12-hour dark/light schedule. Ageing flies were kept 25/vial. They received organic fly food M without Tegosept from Lab Express. They were flipped weekly onto fresh food. We used the w^1118^ line (Bloomington *Drosophila* Stock Center stock #3605) and the Harwich line (Daniel Hartl at Harvard).

### Backcrossing

Cross 1 was between a virgin female Harwick and a male w^1118^ fly. Crosses 2-10 were between a red-eyed virgin female from the previous cross and a w^1118^ male (figure S1). Flies from cross 10 (1:1:1:1 ratio of red-eyed males, red-eyed females, white-eyed males, and white-eyed females) were used in all phenotyping assays except for the hatching time assay, immune, and oxidative stress assays; homozygous lines were established for these assays due to inability to differentiate w^+^ and w^-^ embryos and larvae. To establish the homozygous *white* line, white-eyed virgin females and white-eyed males from cross 10 were crossed (cross 11W). To establish the homozygous red-eyed line, red-eyed virgin females and red-eyed males from cross 10 were crossed (cross 11R). Red-eyed females and red-eyed males from the cross were individually paired in vials (cross 12R) and vials containing any white-eyed offspring were removed; remaining vials with at least 6 red-eyed males (.5^6^x100% = 1.5625% chance of mother being heterozygotic) were used to expand the line. The resulting generations were monitored for white-eyed flies.

### Rapid Iterative Negative Geotaxis (RING) assay

24 hours before the assay, flies were separated into vials of 10 per replicate. At the time of the assay, flies were flipped into empty plastic vials and placed into the Locomotor Activity Monitor (LAM25, Trikinetics) which was attached to a standard multi-tube vortexer (Talboys). Every 30 seconds, flies were shaken to the bottom of the tubes for 3 seconds at the 4^th^ intensity setting and subsequently allowed to climb upwards. The assay was allowed to run for 20 consecutive rounds of shakes (10 minutes). Average number of climbs per fly over 20 consecutive shakes was calculated for each 5 second interval and values for the 10, 15, 20, and 25 second intervals were added to the values from previous intervals to get total number of climbs per fly at the 5, 10, 15, 20, and 25 second mark post-shake.

### Activity assay

Flies were individually placed into a Trikinetics DAM2 monitor and allowed to acclimate for roughly 10 hours before the start of the assay period. The flies’ movement was measured in one-minute intervals for the next 48 hours. The *Drosophila* sleep and circadian analysis MATLAB program (SCAMP) was used for initial analysis, and principal component analysis was done in R^37^.

### Social spacing assay

Social spacing was measured using a modified version of previously described methods^2^. All testing was conducted at the same time of day, between ZT5 and ZT9. Flies were briefly anesthetized with CO_2_ and separated into groups of 20 before being placed in a vertical triangular arena (height = 13”, base = 13”). Once all flies had recovered from anesthesia, they were gently tapped to the bottom of the arena to ensure a uniform starting position. After 20 minutes and the flies fully stabilized into their preferred social spacing, an image was captured for analysis. The position of each fly was identified using segmentation, and the distance to its nearest neighbor (NND) was calculated. Median values of the NND of all 20 flies within the triangle were plotted and analyzed using Kruskal-Wallis statistical testing.

### Immune assay

Innate immune response was assessed using procedures adapted from previously defined methods^55^. Flies were mated on a glucose-yeast diet (15 g glucose/L), and their offspring were aged until they reached the L3 wandering larval stage. To facilitate larval collection, a 4M sucrose solution (150 g sucrose/L) was added to each vial, allowing any remaining larvae to float into the solution. Male and female larvae were then separated, and each individual was gently poked with a 0.25 mm tungsten needle in the cranial third region. The larvae were subsequently transferred to a 96-well plate containing the 4M sucrose solution. After 30 minutes, they were examined for the presence of a black dot, indicating an immune response marked by melanization.

### Lipid Peroxidation

Quantification of lipid peroxidation using BODIPY 581/591 C11 (Invitrogen) was performed as previously described in literature^56^. Briefly, embryos or newly hatched L1 larvae were collected and washed in 1xTSS (0.4% w/v NaCl, 0.03% v/v Triton-X) before being separated into replicates of 10 individuals each in 15 μL of 1xTSS. 63 μL 2xTSS, 60 μL BODIPY solution (1uL of 5 μm BODIPY 581/591 C11 lipid peroxidation sensor in DMSO per 1 mL ddH2O), and approximately 10-20 Lysing Matrix D beads (MP Biomedicals) were added to each sample. Samples were homogenized using a FastPrep-24 Machine at 6m/sec for 30 seconds, then spun down in a microcentrifuge at 6,000 rpm for 1 minute before incubating 30 minutes in the dark at room temperature. Blank samples consisted of 63 μL 2xTSS, 60 μL BODIPY solution, and 15 μL 1XTSS. 100 μL of each sample was transferred to a 384 well plate and read at 485nm and 561 nm on a Synergy Neo2 plate reader. Values were calculated by subtracting the blank values from the sample values, then dividing the 485nm reading by the 591nm reading.

### Triglyceride Assay

Triglyceride content was measured via colorimetry in 8-day old males and mated females as previously described in literature^57^. Each sex/eye color condition had 5 replicates of 10 flies each. The total weight of flies per replicate was recorded before flash-freezing in liquid nitrogen. Tubes were placed on ice, and approximately 0.2g of Lysing Matrix D beads (MP Biomedicals) and 400 μL PBST (0.05% Tween) were added. Samples were homogenized using a FastPrep-24 Machine at 6 m/sec for 20 seconds, then briefly spun down in a microcentrifuge before incubating at 70°C for 10 minutes. Samples were homogenized again at the same settings, then spun down at 10,000 rpm for 5 minutes at room temperature. 40 μL of supernatant from each sample was transferred to a 96-well plate, with 2 technical replicates per sample. 200 μL of triglyceride working reagent (Sigma) was added to each replicate, and 2 wells of 240 μL blank reagent were added to the plate. The plate was sealed and incubated in a 37°C water bath for 10 minutes. The plates were then spun at 4,000 rpm for 5 minutes at room temperature, and 100 μL of each replicate was transferred to a new plate to be read on a Synergy Neo2 plate reader at 540 nm.

### Starvation Resistance Assay

8-day old adult male and mated female flies were separated under mild CO_2_ anesthesia and transferred into vials containing 3 mL of solidified agar media. The assay was set up with five replicates per sex and eye color and 20 flies per replicate. The number of dead flies was recorded every 2 hours from 6 AM to 10 PM daily until all flies had died. The average starvation resistance was calculated in hours as the grand mean lifespan across the five replicates.

### Fecundity and Fertility Assay

The fecundity (unmated females) and fertility (mated females) assay was conducted to quantify egg output across three successive age groups: age 11 (days 10-12), age 21 (days 20-22), and age 31 (age 30-32). The assay used 20 females per replicate with 6 replicates per group over 3 days, with each day counted separately. Virgin females were collected to assess fecundity, while mated females were collected to assess fertility. For fertility assays, non-virgin females were mated 24 hours before the assay. Plates were flipped and eggs were counted every 12 hours.

### Longevity Assay

Newly hatched flies from cross 10 were collected under mild CO_2_ anesthesia and separated into vials based on sex and eye color, with 10 flies per vial and 10 replicates per sex/eye color combination. Vials were checked daily for fly deaths and flipped onto new food every Tuesday and Friday. The assay continued until all flies had died. Lifespan was recorded as the number of days post-emergence. Each replicate value represents the average lifespan of the 10 flies in a vial.

### RNA isolation

RNA was isolated from 1-week old unmated males and females. For each sample, 10 heads were removed under mild CO_2_ anesthesia and flash-frozen in liquid nitrogen. 500 μL of Trizol (Invitrogen) and approximately 0.2g of Lysing Matrix D beads (MP Biomedicals) were added to each sample, and samples were homogenized using a FastPrep-24 Machine at 6 m/sec for 30 seconds 3 times with a 5-minute rest on ice in between homogenizations. Samples were vortexed for 15 seconds and incubated at room temperature for 5 minutes before adding 100 μL of chloroform, vortexing another 15 seconds, and incubating at room temperature for 5 minutes. Samples were centrifuged at 12,000xg for 15 minutes at 4°C. 200 μL of the upper aqueous layer was transferred to a new tube, where 1 μL of GlycoBlue and 225 μL ice-cold isopropanol were added. Samples were incubated 10 minutes at room temperature before centrifuging at 12,000xg for 15 minutes at 4°C. The supernatant was removed, and the pellet was washed twice with 1 mL 75% ethanol and then spun at 7,500xg for 5 minutes at 4°C to remove the 75% ethanol. Samples were allowed to dry for 10 minutes before rehydrating with 15 μL nuclease-free water and incubating at 60°C for 5 minutes.

### RNA sequencing

Libraries were prepared using 500 ng RNA and the Kapa mRNA Hyperprep Kit (Roche) according to the manufacturer’s protocol. Samples were sequenced using an Element AVITI to a depth of 30 M reads and a run length of 2×75bp.

### Differential expression, gene set enrichment, and over representation analysis

Low-quality read ends were trimmed, followed by adapter sequence removal using Trimgalore v0.6.1^58^. The --quality and –length parameters were set to 20. Trimmed reads were then aligned to the r6.60 version of the *Drosophila* genome using STAR v2.5.4b^59^ with default parameters. ReadsPerGene.out.tab files were imported into R v4.4.1^60^ to obtain a counts table. DESeq2 v1.44.0^61^ was then run to detect differentially expressed genes (supplemental file 1). Gene set enrichment was performed in R using GSEA 4.3.3^62^, gene set size filters (min=10, max=500), weighted enrichment statistics and Signal2Noise gene ranking. .gmt files for enrichment analsyis were generated using code presented in “reference code to generate gmt file”. The results of the enrichment analysis were displayed in a network plot created using Cytoscape 3.10.1 (cite PubMed ID: 14597658.), plotted are pathways with q-value<0.1 and overlap similarity coefficient >0.1 (figure 4C). Annotation clusters were identified using AutoAnnotate^64^. Results of the GSEA and Cluster analysis can be found in the supplement (tables 1-4). Counts and statistical analyses for up and downregulated (adjusted p-value < 0.05) gene overlaps between males and females were done in R using the GeneOverlap package v1.40.0^63^ (figure 4B, supplemental file 2). Differentially expressed genes with an adjusted p-value cutoff of < 0.05 were used to perform the overrepresentation analysis in WebGestalt (2024 release)^65^ (figure 4D-G, supplemental files 3 and 4).

